# How many ecological niches are defined by the superabundant marine microbe *Prochlorococcus?*

**DOI:** 10.1101/2022.11.29.517206

**Authors:** Miriam Miyagi, Maike Morrison, Mark Kirkpatrick

## Abstract

Determining the identities, frequencies, and memberships of ecotypes in *Prochlorococcus* and other superabundant microbes (SAMs) is essential to studies of their evolution and ecology. This is challenging, however, because the extremely large population sizes of SAMs likely cause violations of foundational assumptions made by standard methods used in molecular evolution and phylogenetics. Here we present a tree-free likelihood method to identify ecotypes, which we define as populations with genome sequences whose high similarity is maintained by purifying selection. We applied the method to 96 genomes of the superabundant marine cyanobacterium *Prochlorococcus* and find that this sample is comprised of about 24 ecotypes, substantially more than the five major ecotypes that are generally recognized. The method presented here may prove useful with other superabundant microbes.

## INTRODUCTION

With densities of up to 10^6^/ml of seawater and a global census size of some 10^27^ cells, the cyanobacterium *Prochlorococcus marinus* is the Earth’s most abundant photosynthetic organism (1). It is also among the most ecologically important: these microbes are responsible for some 10% of atmosphere’s oxygen (2).

The taxon *Prochlorococcus* encompasses multiple ecotypes that are distinguished by physiology, genomic features, and the environments in which they live (e.g. (3-11)). The number, frequencies, and memberships of these ecotypes, however, is not well understood. This situation is partly the result of the differing criteria used to delimit the ecotypes and estimate their phylogenetic relations: nucleotide or amino acid identity (*e.g.* (3, 7, 12)), physiology (13, 14), likelihood (15-17), neighbor joining (7, 18), parsimony (19), and the frequency of recombination (20).

Having accurate definitions for the ecotypes is important for several reasons. It is of great interest to predict how *Prochlorococcus* and other superabundant photosynthetic microbes will respond to climate change. The accuracy of current models (1, 21-23) might be increased by explicitly recognizing the ecological and physiological substructure of *Prochlorococcus* (14). Understanding the molecular evolution of *Prochlorococcus* depends on appropriate definitions of genetic populations, but studies have used widely differing ones (12, 19, 24, 25). Defining ecotypes in superabundant microbes presents several novel challenges. Current methods in molecular evolution and phylogeny assume that a certain class of genomic sites (*e*.*g*. synonymous site) evolve as selectively neutral, and that each nucleotide variant descend from of a unique mutation. If there are selectively neutral mutations occurring at some genomic sites, however, the vast coalescence times implied by the huge population size suggest that these sites may be mutationally saturated, and all trace of phylogenetic history has been erased. There is further the question of whether effectively neutral mutations (that is, with *N*_*e*_ *s* << 1) even exist in the genome of an organism with the population size of *Prochlorococcus* (18). A recent study suggests *N*_*e*_ may in fact be only on the order of 10^7^, or the number of cells than can be found in 10 ml of seawater (26). That conclusion, however, was based on the assumption that some sites in the genome are selectively neutral. Clearly there is need for molecular tools that are appropriate to the biology of SAMs.

Some intuition about the implications of the extreme population size is gained from Figure 1. It shows how nucleotide diversity (π) varies with population size, selection, and drift. These are results from a toy model that is described in Supplemental Information 1. Although it is far too simplified to rely on for quantitative results, the qualitative outcomes are informative. For most of life on Earth, the product of population size and the per-base mutation rate (*N*_e_ μ) is much smaller than 1. In that realm, the evolution of a mutation will either be largely dominated by mutation and drift (if *N*_*e*_ *s* << 1) or by mutation and selection (if *N*_*e*_ *s* >> 1). Standard methods for inferring the genetic boundaries of species and their phylogenetic relations rely on mutations in the first category (*e*.*g*. refs. (27, 28)). But when *N*_e_ μ is much greater than 1, sites whose evolution is dominated by mutation and drift no longer occur. Instead, allele frequencies are either determined by mutation rates alone (if *N*_e_ *s* is sufficiently small) or a balance between mutation and selection (if not). In either event, we do not have the class of mutations required by standard methods that aim to delimit populations and species, and to estimate the phylogenetic relations between them.

**Figure 1.**
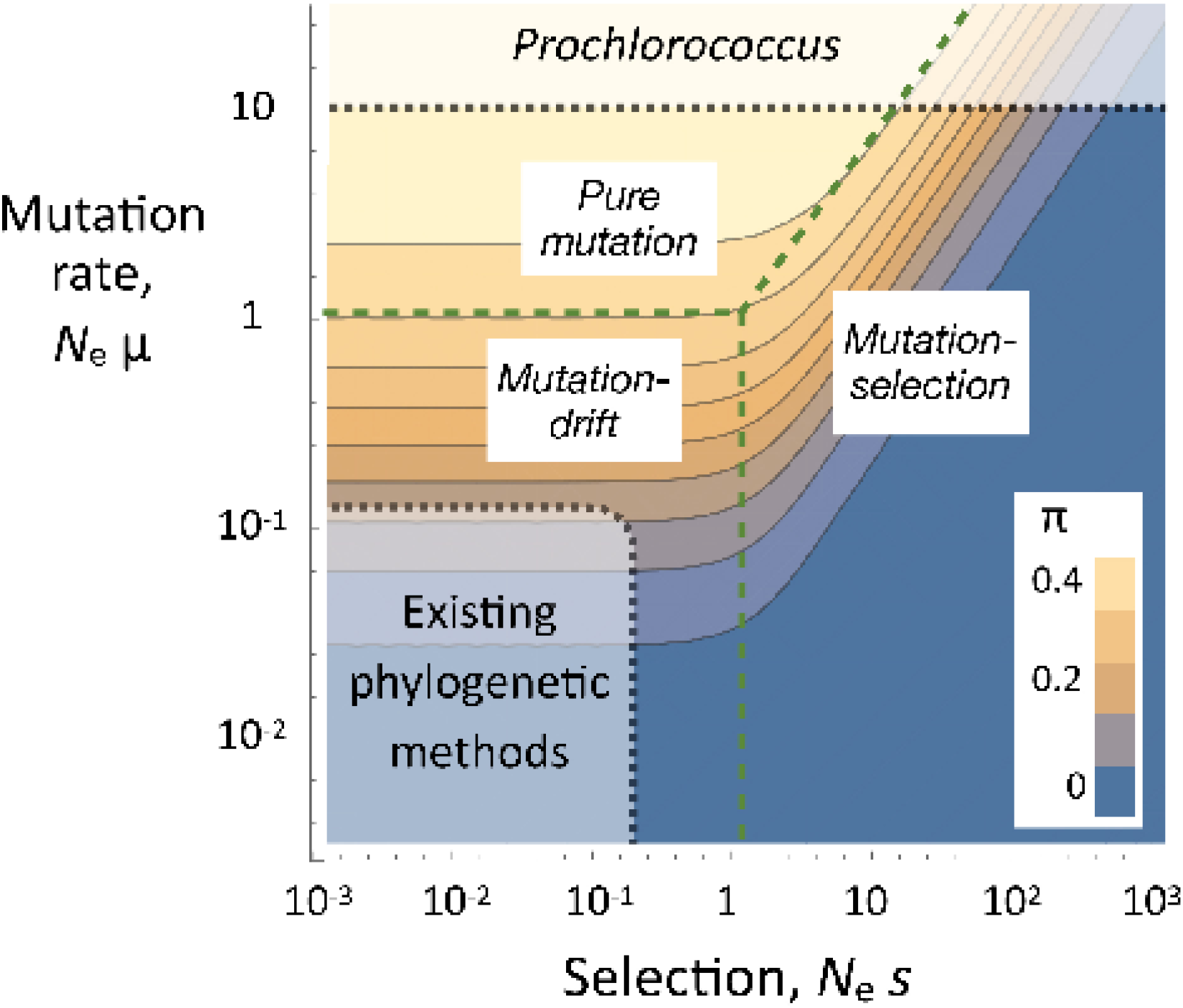
The dependance of nucleotide diversity (π) on the scaled strength of selection (*N*_e_s) and mutation rate (*N*_e_μ). The results are based on the toy model described in detail in Supplemental Information 1. The population size of *Prochlorococcus* is so vast that it may lie outside the region of parameter space assumed by existing phylogenetic methods.

In this paper we introduce a new model for delimiting ecotypes in superabundant microbes that we call *TreeFree*. We use it to analyze 96 whole genome sequences and 101 internal transcribed spacer (ITS) sequences sampled from *Prochlorococcus* by Kashtan *et al*. (18). The basis for our approach is an operational definition of an ecotype as a population whose individuals experience such similar selection pressures that they have very similar genomes and (by implication) ecological function. This view of ecotypes has some similarity to Cohan’s definition (29), but ours does not require periodic selective sweeps to homogenize the genetic variation within ecotypes since that can be accomplished by purifying selection alone. The core assumption is that each ecotype is characterized by a reference genome sequence, and the rare departures from that sequence result from deleterious mutation (and sequencing errors). Under this hypothesis, any variation maintained by some form of balancing selection in effect results in multiple ecotypes that coexist by some form of niche partitioning. Our method uses a likelihood approach to estimate the number of ecotypes and their frequencies, and there is an explicit model for the sources of genetic variation within ecotypes (mutation and sequencing error). No attempt is made to estimate the phylogenetic relations between the ecotypes: the method is tree-free. The reference sequences for all the ecotypes appear in the likelihood function, but since those are not our main concern we average over uncertainties in those sequences using Markov Chain Monte Carlo (MCMC) (30).

We note that the term “ecotype” has been used in ways that differ from our definition by researchers working on *Prochlorococcus* and other microbes. In this paper, an ecotype refers to a group of cells that our analyses suggest belong to the same genetic unit (as described above). We use “clade” to refer to the groups of genomes recognized by Kashtan *et al*. (18). (Later we will introduce “population” to refer to other levels of clustering.)

To learn how results from *TreeFree* compare with those from conventional methods used to infer species boundaries with genomic data, we also analyzed the Kashtan data using Bayesian Phylogenetics and Phylogeography (BPP), a Bayesian method for delimiting species (or in our case, ecotypes) using the multispecies coalescent model (31, 32). As with other methods in molecular phylogenetics, BPP makes assumptions we suspect may be violated by *Prochlorococcus*. Notably, BPP assumes that genomic sites that differ between species (or ecotypes) are not mutationally saturated, and that there is strong recombination between sites.

Results from our new method suggest that there are many more ecotypes than are generally recognized by the community of *Prochlorococcus* workers and by BPP. We estimate that this sample of 96 genomes comprises about 24 ecotypes. Our method is many times more computationally efficient than BBP, and it can accommodate sequences of 1 Mb or more. This tool may be useful for exploring the ecological diversity in other superabundant microbes.

## METHODS

### The data

This study was inspired by Kashtan *et al*. (18), who collected partial genome sequences from 96 *Prochlorococcus* cells that were sampled from 2 ml of seawater at 60 m depth at a site in the mid-Atlantic during three dates in 2008 and 2009. In addition, we analyzed the RNA ITS (549 bp) that was sequenced from those 96 cells and from an additional five cells.

The genomes of *Prochlorococcus* ecotypes differ dramatically in their size and composition due to variation in the presence or absence of “flexible” genes. Since our method relies on sequence alignment, we focused exclusively on the 1 Mb “core” genome that is shared among all ecotypes and consists of about 1 400 genes. There are 307 432 SNPs. Of these, an unusually high fraction (21%) are triallelic or quadallelic, which is not surprising given the presence of multiple ecotypes.

### TreeFree

We call our new method *TreeFree* because it estimates the genetic boundaries of ecotypes without estimating their phylogeny. At the heart of the algorithm is a model that calculates the probability of the observed genome sequences given four sets of parameters: the number and frequencies of ecotypes, the assignment of each genome in the sample to an ecotype, the reference genome sequence for each ecotype, and the frequency of minor alleles at sites within ecotypes that result from deleterious mutation and sequencing error. Our main interest is in the first of these.

Given this model, one strategy might be to search for the parameters that maximize the likelihood (that is, the maximum likelihood estimates). Unfortunately, that strategy is not practical because of the very large number of parameters, notably the reference genome sequences for all the ecotypes. For example, a statistical model for a dataset with 10 ecotypes with genomes that have 10^5^ SNPs has 10^6^ parameters to estimate. We therefore use Markov Chain Monte Carlo to sample the parameter space and obtain posterior probability densities for the parameters of interest. A technical challenge here is that we need to compare the likelihoods for models that include different numbers of ecotypes and therefore different numbers of parameters.

The following two sections outline the likelihood model and the MCMC algorithm. Details are given in Supplemental Information 2.

#### The likelihood model

The essence of the likelihood model is simple. For each genomic sequence in the sample, we consider the probability that it belongs to each proposed ecotype. For each of those ecotypes, we calculate the probability of the observed sequence, which is determined by the numbers of sites at which that sequence does and does not agree with the reference sequence for that ecotype. The probabilities for membership in each ecotype are added together to give the total likelihood for that sequence. The likelihoods for each sequence in the sample are multiplied together to arrive at the likelihood for all of the data, given the parameters. This last step assumes that the individual genomes are independent samples.

We greatly simplify this model by making two strong assumptions. First, we assume that within an ecotype, the minor allele at each site has the same frequency q. This assumption is plausible if purifying selection is sufficiently strong that the great majority of genomic sites fall into the mutation-selection domain shown in Figure 1 and if most minor alleles result from sequencing error rather than deleterious mutation. The latter assumption seems plausible since the sequencing error in these data is estimated to be 10^−4^ per base pair (18) while the spontaneous mutation rate in *Prochlorococcus* is on the order of 10^−10^ per base pair (26, 33).

Under these assumptions, the likelihood of the data is

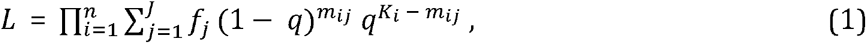

where *n* is the number of sequences in the sample, *J* is the proposed number of ecotypes, *f*_*j*_ is the frequency of ecotype *j, K*_*i*_ is the total number of SNPs in sequence *i*, and *m*_*ij*_ is the total number of matches across all genomic sites between the alleles in sequence *i* and those in the reference genome of ecotype *j*. Further details are given in Supplemental Information 2.

#### The MCMC implementation

We infer the ecotype structure in a way similar to the implementation of *structure* (34). For a given number of ecotypes, we alternate between Metropolis-Hastings steps in order to optimize the vector of ecotype frequencies and Gibbs steps to estimate the reference sequences for all the ecotypes (see (35)).

We initiate that algorithm with *J* (the number of ecotypes) equal to *n* (the sample size), so that each genome is initially assigned to a different ecotype. We then decrement the number of ecotypes by one, removing the ecotype that causes the smallest change in the likelihood when it is omitted. After iterating this process down to a single ecotype, we take the maximum likelihood achieved within each Gibbs step. This likelihood is compared with the maximum likelihood reached in the previous Gibbs step using a likelihood ratio test. These steps are repeated until only a single ecotype remains. We then count the number of times that a Gibbs step results in a significant decrease in the likelihood (at *p* < 0.05). This number is our estimate for the number of ecotypes in the sample.

The logic behind this algorithm is as follows. Whenever removing an ecotype causes a significant drop in the likelihood, we expect that this potential ecotype is in fact a real ecotype. Conversely, if removing an ecotype does not cause the likelihood to drop significantly, we interpret reject that potential ecotype as being a real one.

We found that using subsets of the genome produces smaller estimates of the numbers of ecotypes. This behavior is expected because the sensitivity of the likelihood scales with the length of the sequences.

#### Data analyzed

We found it was not feasible to run TreeFree on the full sequences. We therefore analyzed the first 1% of the genome, and the first 10% of the genome.Comparisons between these two analyses show how sensitive our method is to the amount of data.

### BPP

We compared the results from our method with those obtained from Bayesian Phylogenetics and Phylogeography (BPP), a Bayesian method for delimiting species (or in our case, ecotypes) using the multispecies coalescent model (31, 32). We applied this method to the genomes sequenced from single cells of *Prochlorococcus* by Kashtan *et al*. (18). Ninety of these genomes come from what they refer to as ecotype cN2.

The BPP analysis proceeds as follows:

1. Each genome is assigned to a small “population” that is *a priori* assumed to belong to only one ecotype. A rooted “guide tree” is provided that gives an initial phylogeny for these populations. For this purpose, we used the phylogeny proposed by ref. (18) (see Fig. 2).
2. BPP uses a Markov Chain Monte Carlo (MCMC) algorithm that considers jumps to different guide tree topologies. A reversible-jump MCMC algorithm considers changes to ecotype delimitations by merging and splitting tips of the guide tree. This process iterates many times.
3. BPP outputs posterior probability distributions for several quantities, notably the total number of ecotypes and the assignments of each genotype to an ecotype.

**Figure 2.**
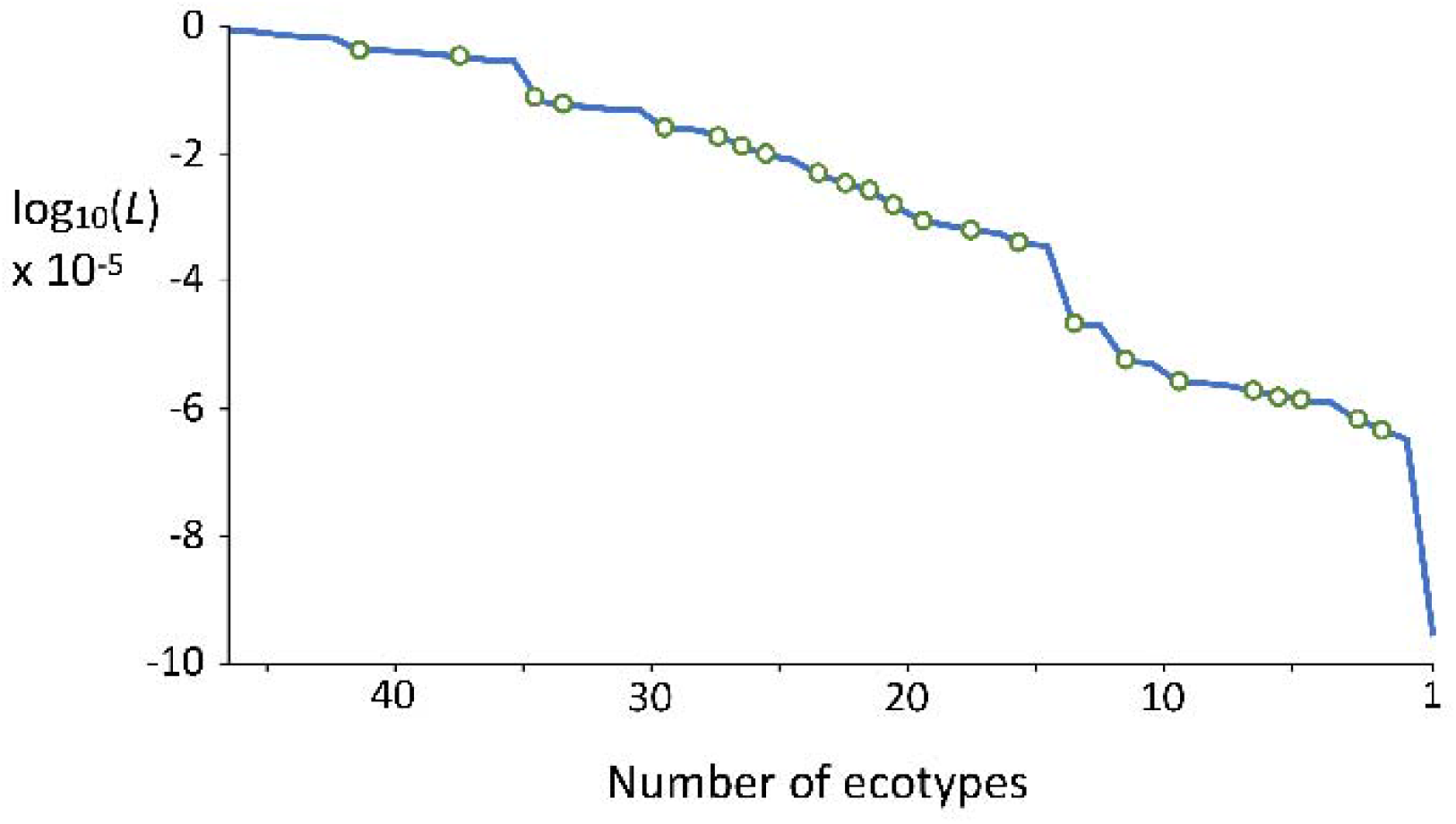
The log likelihood of ecotype partitions calculated by *TreeFree* using 10% of the genome. Moving to the right, in each Gibbs step the number of ecotypes is decreased by one, resulting in a decrease in the likelihood. Steps in which the decrease is significant (*p* < 0.05 by a likelihood ratio test) are indicated by the circles.

BPP is unable to run using the entire whole genome data set. We therefore ran it on three subsets of the sequences: about 10% of the core genome (163 kb, *n* = 96), about 0.1% of the core genome (1.63 kb, *n* = 96), and the rRNA ITS sequences (549 bp, *n* = 101 sequences). Below we report the results from 12 distinct analyses that differ in the dataset used (a proportion of the core genome or the entire ITS region), the sequences that were included, and the assignment of sequences to populations (see Supplemental Information Table SI 3.1).

For many of our analyses, we focused on one or more ecotypes, initiating BPP with two or more prior populations from each ecotype. We then observed if BPP assigned these prior populations to their own ecotypes or merged several populations into the same ecotype. A more thorough discussion of our BPP implementation is included in the Supplemental Information 3, and details of 12 selected analyses are given in Supplemental Information Tables SI 3.1 and SI 3.2.

## RESULTS

### Results from *TreeFree*

Both *TreeFree* and BPP are Bayesian methods, so rather than providing single point estimates of parameters they return the probabilities associated with all possible outcomes. For brevity, in the text we will refer to the result that has the highest posterior probability. As described above, we used *TreeFree* to analyze subsets of 0.1% and 10% of the sites in the core genome from all 96 individuals. Using 0.1% of the sites, the number of ecotypes with the highest posterior probability was about 7, while with 10% of the sites it was about 24 ecotypes. More details of the results are presented in Figure 2 and Supplemental Information 4.

Notably, the 24 ecotypes identified in the larger dataset are not perfect subsets of the 7 ecotypes found using the smaller dataset. This suggests that additional sequence data are required not only to resolve ecotypes on finer scales, but also to determine whether ecotypes have been robustly identified. This outcome is not entirely surprising since smaller subsets of the genome can leave out genes that are critical to ecological differences between the ecotypes.

### BPP Results

We conducted three main categories of analyses. First, we analyzed three subsets of the whole genome sequences: 100%, 10%, and 0.1% (Table 3.1, rows 1-3). Due to the computational limits of BPP, we could not analyze the full genome sequences of all 96 cells at once. Our analysis of the full genomes of nine individuals (four from clade C1, four from C2, and one from cN1-C9) identified three ecotypes which corresponded to Kashtan et al.’s three clades (Table 3.1, row 3). In order to analyze all 96 single-cell sequences at once, we restricted our analysis to either 10% or 0.1% of the full genomes. Here we initiated BPP by dividing each major clade into 2 populations. Both analyses estimated that the sequences belonged to between 13 and 14 ecotypes, with the 10% analysis placing slightly more weight on larger numbers of ecotypes and the 0.1% placing more weight on smaller numbers (Table 3.1, rows 1-2). Second, we analyzed the 549 bp of the ITS rRNA sequences from 101 genomes. We divided the four largest Kashtan clades (C1, C2, C3, and C4) into multiple prior populations, and initiated BPP with the neighbor joining tree estimated by Kashtan *et al*. (2014) (shown in Figure 3). BPP merged the populations within each clade, resulting in 14 ecotypes (Fig 3; Supplemental Table SI 3.1, row 4). These are largely consistent with the ecotypes and clades recognized by Kashtan *et al*., but two of the clades are split into a pair of ecotypes. Second, we used a random guide tree. BPP then merged the populations further, resulting in 10 ecotypes (Supplemental Information Table SI 3.1, row 5).

**Figure 3.**
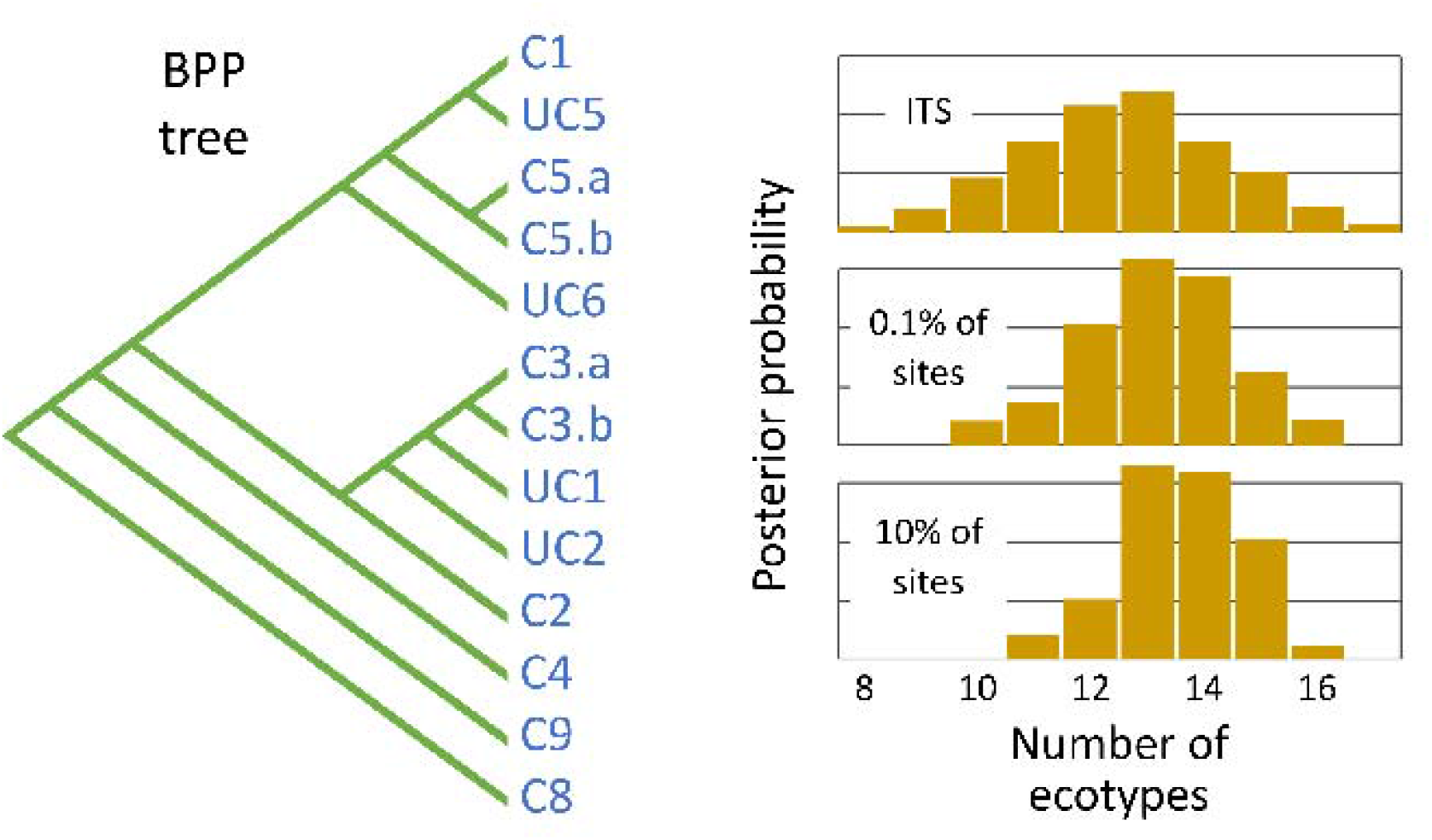
*Left*: The most probable phylogenetic tree estimated by BPP using 10% of the genome sequences. The ecotypes it identified are consistent with the clades identified by Kashtan *et al*. (18) with the exception of clades C3 and C5, which BPP subdivided into two ecotypes. For brevity, Kashtan *et al*.’s clades c9301-C8 and cN1-C9 are shown here as C8 and C9. Ecotypes UC1, UC2, and UC5 are represented by only a single genome. Other ecotypes are represented by between 2 and 53 genomes; ecotype C1 is by far the most abundant. *Right*: The posterior distributions of probabilities for the numbers of ecotypes estimated by BPP using the ITS sequence alone, 0.1% of the genomes, and 10% of the genomes.

Finally, we ran BPP on a reduced number of single-cell ITS sequences. By including fewer individuals in the analysis, we were able to initialize BPP with a larger number of populations, each with fewer individuals. These analyses allowed us to test the extent to which BPP over-split populations. When we assigned each of the 13 individuals in clade C3 to its own initial population, BPP merged all of them into a single ecotype. Similar outcomes obtained with other initial populations, with the exception that one initialization led to multiple ecotypes within clade C1 (Supplemental Information SI 3.1, rows 8 – 12).

Overall, the ecotypes returned by BPP are largely consistent with the clades identified by Kashtan *et al*. (18) using neighbor joining. There are big differences, however, in the topology of the trees estimated by BPP using 10% of the sequences and neighbor joining using the whole genomes (Figure 4).

**Figure 4.**
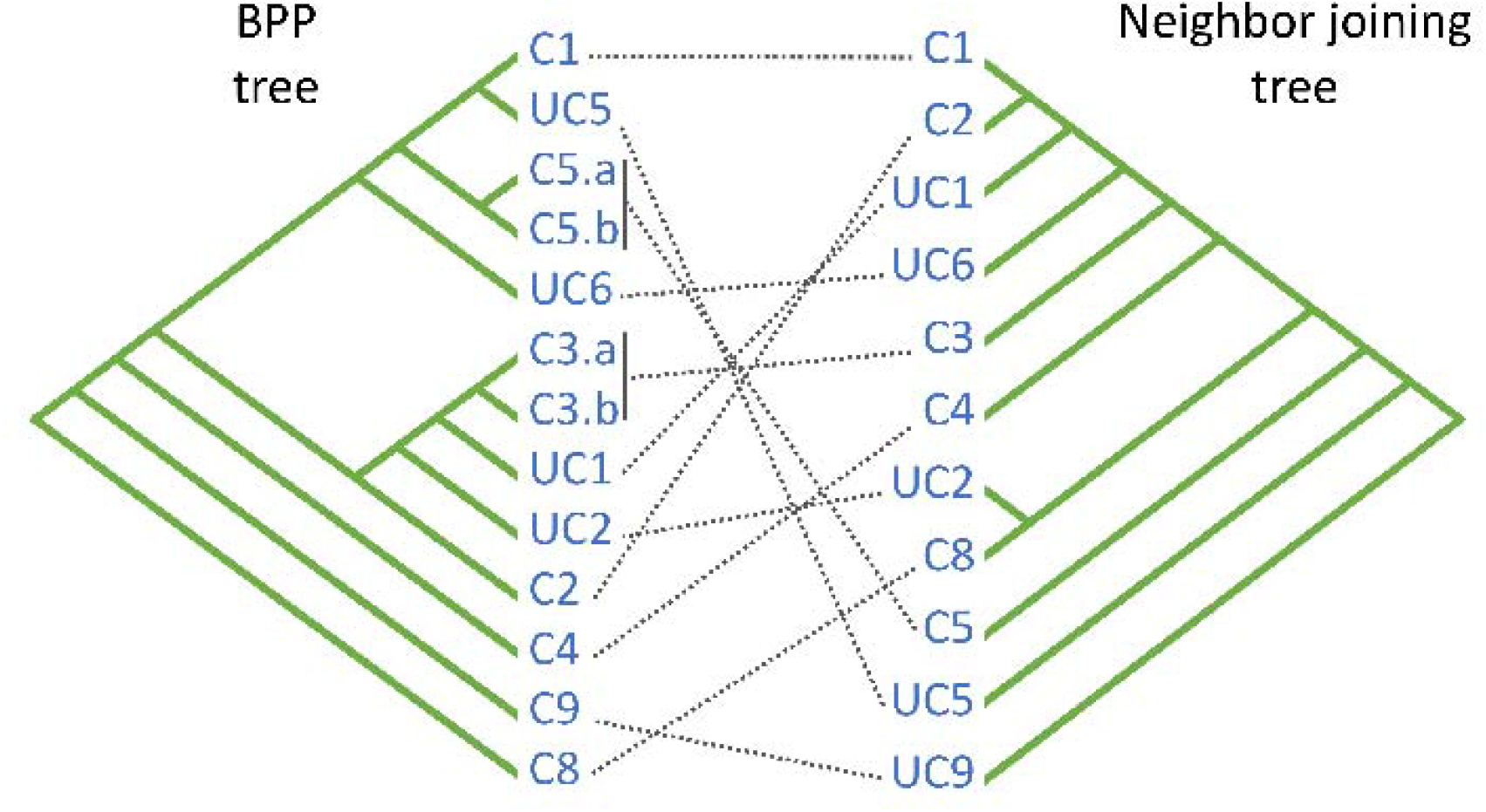
The relationship between the phylogenetic tree estimated by BPP using 10% of the data and the neighbor joining tree estimated by Kashtan *et al*. (18) using the whole genomes.

### Comparing *TreeFree* and BPP

The most important difference between the results from *TreeFree* and BPP is the number of ecotypes estimated. Using 10% of the sequences, *TreeFree* estimates that there are roughly twice as many as does BPP (about 24 *vs*. about 12).

The two methods also disagreed on some of the assignments of the genomes to ecotypes (Figure 5). For example, *TreeFree* subdivided the C1 clade while BPP did not, and *TreeFree* cleanly divided the cN1-C9 and cN1 clades into separate ecotypes, rather than lumping them together. On the other hand, *TreeFree* lumped the UC and C5 clades into a single ecotype, which BPP did not.

**Figure 5.**
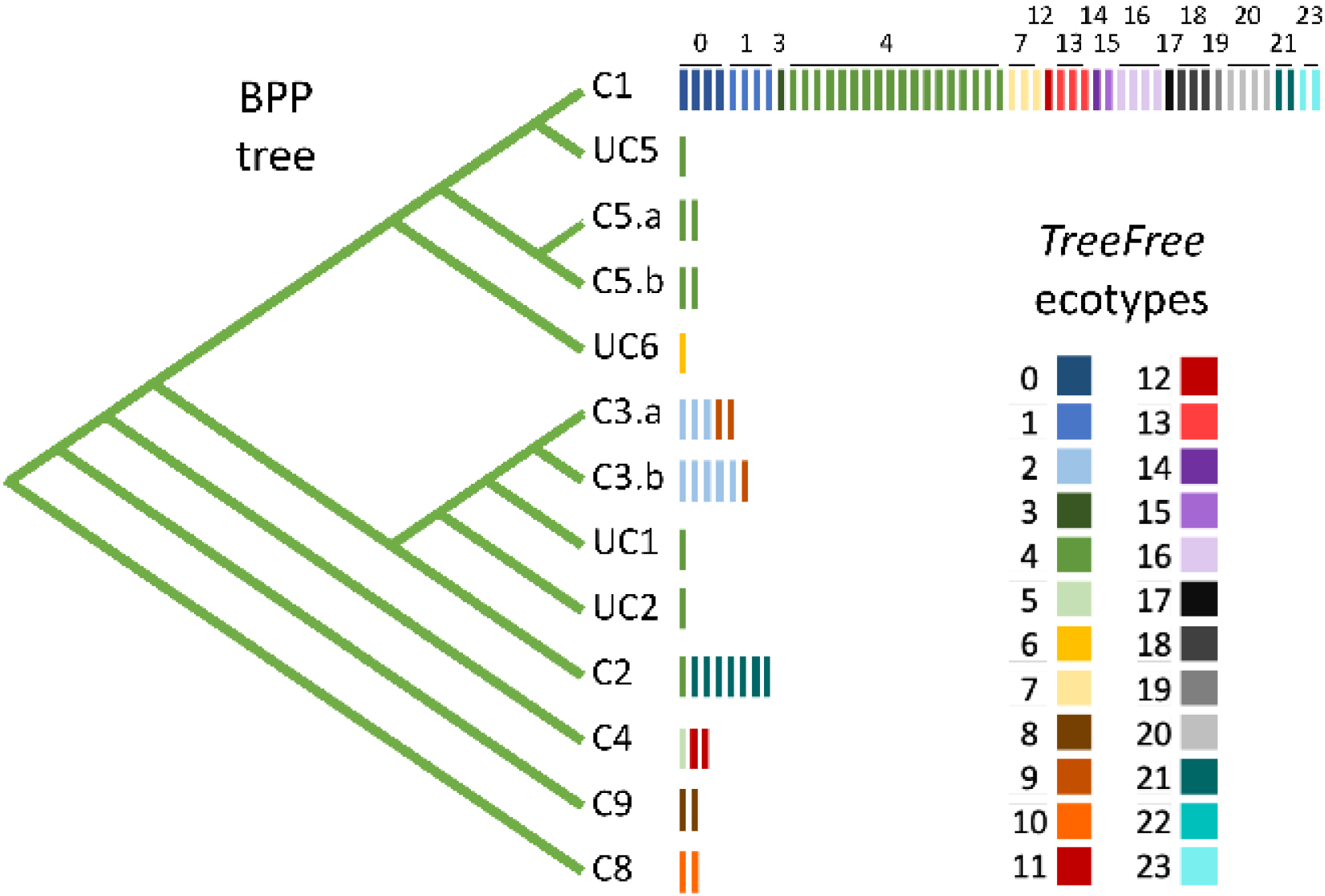
The relationship between the ecotypes estimated by *TreeFree* and BPP using 10% of the sequences. At left are the phylogeny and ecotypes estimated by BPP (see Figs. 3 and 4). To the right of the tips of the tree, each vertical rectangle represents one of the genomes sequenced by Kashtan *et al*. (188), color coded to show to which of the 24 ecotypes they most likely belong according to *TreeFree*.

## DISCUSSION

Our findings suggest that the number of ecotypes in *Prochlorococcus* may be substantially larger than are commonly recognized. Early biochemical work suggested that *Prochlorococcus* had two ecotypes adapted to high and low light (36). The arrival of genome sequences, larger sample size, and additional environmental data lead to the recognition of six ecotypes (37), and many later studies accepted that conclusion. More recent work has subdivided these further. Kashtan *et al*. (2014) analyzed a sample of 1 381 sequences of the ITS. Using a cutoff of 99% sequence identity, they found that depending on the season between 130 and 200 “backbone subpopulations” coexisted in their samples. Further, by subsampling different numbers of those sequences they showed that the true number of these subpopulations was certainly much larger.

Based on a new method called *TreeFree*, our analysis suggests the presence of about 24 ecotypes in the 96 whole genome sequences sampled by Kashtan *et al*. (2014). The method, which was designed to delimit ecotypes using genomic data from superabundant microbes, is based on an explicit statistical model. Our model makes the strong assumption that the effective strength of selection, *N*_*e*_ *s*, is much larger than one at all sites in the genome. While that assumption is plausible in the case of *Prochlorococcus*, we currently have no direct way to test it directly. Our conclusions are therefore provisional until the arrival of new statistical methods that can estimate quantities from patterns of molecular variation in superabundant microbes.

Properly defining ecotypes in *Prochlorococcus* could open up a new field of molecular evolution. The combined census population size estimated for *Prochlorococcus* is so vast that even ecotypes that are quite rare may have population sizes many orders of magnitude larger than those of abundant eukaryotes such as *Drosophila*. As we suggested in the Introduction, this situation could put *Prochlorococcus* in an unexplored region of population genetics parameter space. If *N*_*e*_ μ is much larger than 1 throughout the genome, all sites will be mutationally saturated. That situation could free *Prochlorococcus* of most adaptive constraints. Adaptive sweeps of point mutations cease to occur because every possible mutation occurs many time in each generation, and most adaptation may happen by selection on standing variation (38). If *N*_*e*_ *s* is much larger than one at all sites, then no mutations evolve as if neutral, and genetic drift is virtually banished as an evolutionary force. This situation would represent a strange and fascinating new world for evolutionary genetics.

## ACKNOWLEDGEMENTS

We are very grateful to Colin Walker for assistance with programming. This research was supported by NSF grant DEB-1831730 and NIH grant R01-GM116853 to MK.

## COMPETING INTERESTS

The authors declare they have no competing financial interests.

## DATA AVAILABILITY STATEMENT

All data analyzed in this study is in the public domain and can be located by consulting the references cited in the text. The scripts and code used in these analyses are available at [*a public repository to be specified before publication*].

## Supplemental Information for

## SI 1: Toy model used for Figure 1

Here we calculate the expected molecular diversity, π, at a site evolving under mutation, selection, and drift. The model is highly simplified and not intended to accurately capture the relevant biology. Further, π by no means gives a complete description of a site’s evolution. The point of these calculations is simply to show that sites with the properties assumed by classical phylogenetic methods do not occur when population sizes are so large that *N*_*e*_ μ >> 1.

The model is of a biallelic locus in a haploid population with constant size *N*. Mutation between the alleles is symmetric at rate μ. The relative fitnesses of the alleles are 1 :: 1 + *s*. We assume the classic Wright-Fisher model of drift.

Wright (1, 2) found that the stochastic equilibrium distribution of allele frequencies is

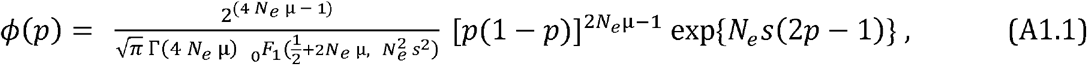

where _0_ *F*_1_ (·, ·)is the regularized confluent hypergeometric function (3). The expected molecular diversity is then

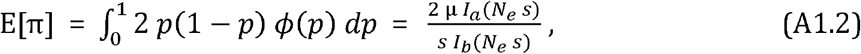

where

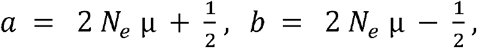

and *I*_*n*_*(z)* is the modified Bessel function of the first kind (3). Figure 1 in the main text is based on Equation (A1.2).

## SI 2: The *TreeFree* algorithm

Our goal is the inference of the frequencies of ecotypes present in our sample and the genome sequences of those ecotypes. As *Prochlorococcus* has an extraordinarily large population size (4), we hypothesize that the frequencies of genotypes within an ecotype are determined by a mutation-selection balance, with most sites in most genomes carrying the reference allele for its ecotype. Accordingly, we designed an algorithm which clusters the genotypes into ecotypes based on sequence similarity in a manner similar to *structure* (5). The algorithm is described in the following, and is summarized with pseudocode in the last section of this appendix.

As in the main text, we use the following notation to describe our implementation:

**X** Data matrix of genome sequences, with *X*_*ik*_ equal to the allele observed at the *k*^th^ site in the *i*^th^ sequence.

***J*** Number of ecotypes in the model

**G** Reference sequences for the ecotypes, with *G*_*jk*_ equal to the allele at the *k*^th^ site in the *j*^th^ ecotype

**f** Vector of estimated ecotype frequencies, with *f*_*j*_ equal to the frequency of the *j*^th^ ecotype

*K* Total number of SNPs in the sample

*m*_*ij*_ Number of sites at which the allele at site *i* and ecotype *j*

*q* Minor allele frequency at all sites

We begin by assuming that each genotype in the sample comes from a different ecotype, and that the reference genome sequence for that ecotype is exactly equal to the sequence of that genotype. (This is the value of **G** with the highest likelihood.) We then decrease the number of ecotypes (*J*) to force multiple genotypes to be clustered within ecotypes, then use an iterative method to approximate the highest likelihood value of **G** given *J*. This scheme is composed of two alternating step. First, a Metropolis-Hastings MCMC step is used to adjust the frequencies of the ecotypes (the *f*_*j*_). Second, a Gibbs MCMC step updates the ecotype genome sequences **G**, conditioned on their frequencies. These steps are described in more detail in the following section.

### S2.1 Metropolis-Hastings MCMC

We use the Metropolis-Hastings MCMC algorithm to explore the space of possible frequency vectors. We start with a vector **f**^(0)^ in which all ecotypes are equally frequent (i.e., the entries of **f**(0) are all equal to 1/*N*). Next we generate a sequence of frequency vectors **f**(^1)^, **f**^(2)^, …, **f**^(*n*)^ using the following rules. Per the Metropolis-Hastings algorithm, we pick a proposed vector f^(*n*+1)^ near to **f**^(*n*),^ then decide whether or not to accept or reject the proposal with probability proportional to the product of the ratio of likelihoods and the probability of sampling one **f** vector given the other. Formally, holding **G** (the genome sequencies for the ecotypes) fixed, we compute:

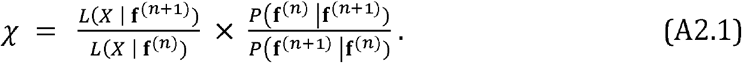

In this equation the likelihood terms are the same as in Equation (1) of the main text, and *P* (**f**^(*n*+1)^ | **f**^(*n*)^)is the probability that we accept the proposed frequency vector **f**^(*n*+1)^. We sample these proposals from a Dirichlet distribution:

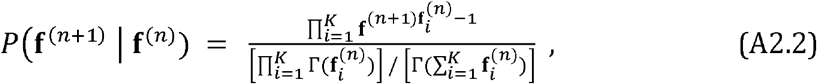

where Γ(·) is the gamma function. We accept a proposed frequency vector **f**(*n*+1) with probability *χ*, and retain the current vector **f**^(*n*)^ with probability (1 − *χ*).

By initializing our MCMC sampler with the value of **G** that maximizes the ikelihood, we minimize the burn-in phase phase for each following value of *J*. To encourage the sampler to sample away from vectors with zero-valued entries, we bounded sampled values from below at a frequency of one individual per million, well below our expected resolution given our sample size.

### S2.2 Gibbs MCMC

To optimize the matrix of reference sequences, **G**, we will use a different MCMC algorithm, the Gibbs sampler. It is well suited for dealing with the categorical nature of the ecotype genome sequences.

We sample the elements of **G** one at a time. For each element, we calculate the likelihood for all four possible bases, and choose among these proposals with probabilities proportional to their likelihoods.

To minimize the numerical burden, we observe that the likelihood (see Equation 1) can be written as:

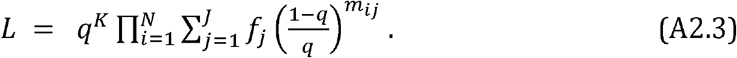

This formulation is convenient because *m*_*ij*_ can only change by one when a base in a reference sequence is changed. We can further reduce computation by fixing the ecotype *j*. Then the likelihoods for all possible alleles at all the sites in that ecotype are given by

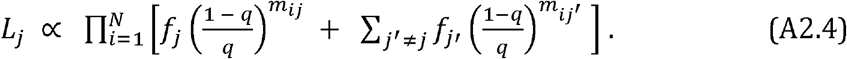

This is useful because the second of the two terms inside the square brackets is constant with *j* fixed. Consequently, that term can be calculated once and used for all the sites within ecotype *j*.

### S2.3 Transitions Between Models

After each set of M-H and Gibbs MCMC steps, we decrement *J* to force clustering of the samples into fewer ecotypes. To define the starting point for the next set of MCMC steps, we remove the ecotype which has the smallest effect on the total likelihood when we reapportion its frequency proportionally to the inverse of the Hamming distance between that ecotype and the remaining ecotypes. That is, to remove ecotype *j* we set

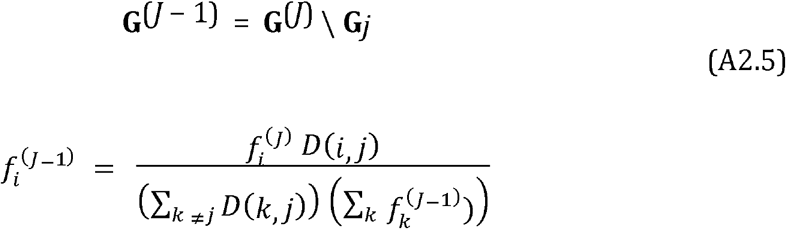

where *D*(*i, j*) is the Hamming distance between ecotypes *i* and *j*. We calculated the likelihood with each ecotype removed, and finally removed the ecotype which resulted in the smallest change to the likelihood.

### S2.4 Estimating the number of ecotypes

Our parameter space is too rich to use standard information criterion tests such as AIC and BIC to choose between alternative estimates of **f** and **G**. We therefore use a simple likelihood ratio test. At each Gibbs step, we found the maximum likelihood among all of the Metropolis-Hastings steps. We then compared these likelihoods among successive Gibbs steps using a likelihood ratio test a difference in the number of parameters equal to the sequence length. Each step resulting in a significant drop in the likelihood (at *p* < 0.05) indicates that a true ecotype has been removed.

The rational for this procedure is as follows. Consider when there are in fact *J* “true” ecotypes, but we are at the Gibbs step with *J* + 1 potential ecotypes. In that case, one of the potential ecotypes is comprised of individuals that in fact belong to one of the *J* true ecotypes. We then expect that its reference sequence will be very similar to that true ecotype. Consequently, assigning the individuals in that potential ecotypes to its true ecotype will result in a small and insignificant drop in likelihood. Conversely, when a Gibbs step removes a true ecotype, we expect the drop in likelihood to be significant. Thus the number of Gibbs steps that result in significant drops in likelihood provides an estimate of the number of real ecotypes in the sample. This sequential pruning of “centers” of potential ecotype is analogous to the “mean shift” procedure that is widely used in pattern recognition (6).

### S2.5 The TreeFree algorithm

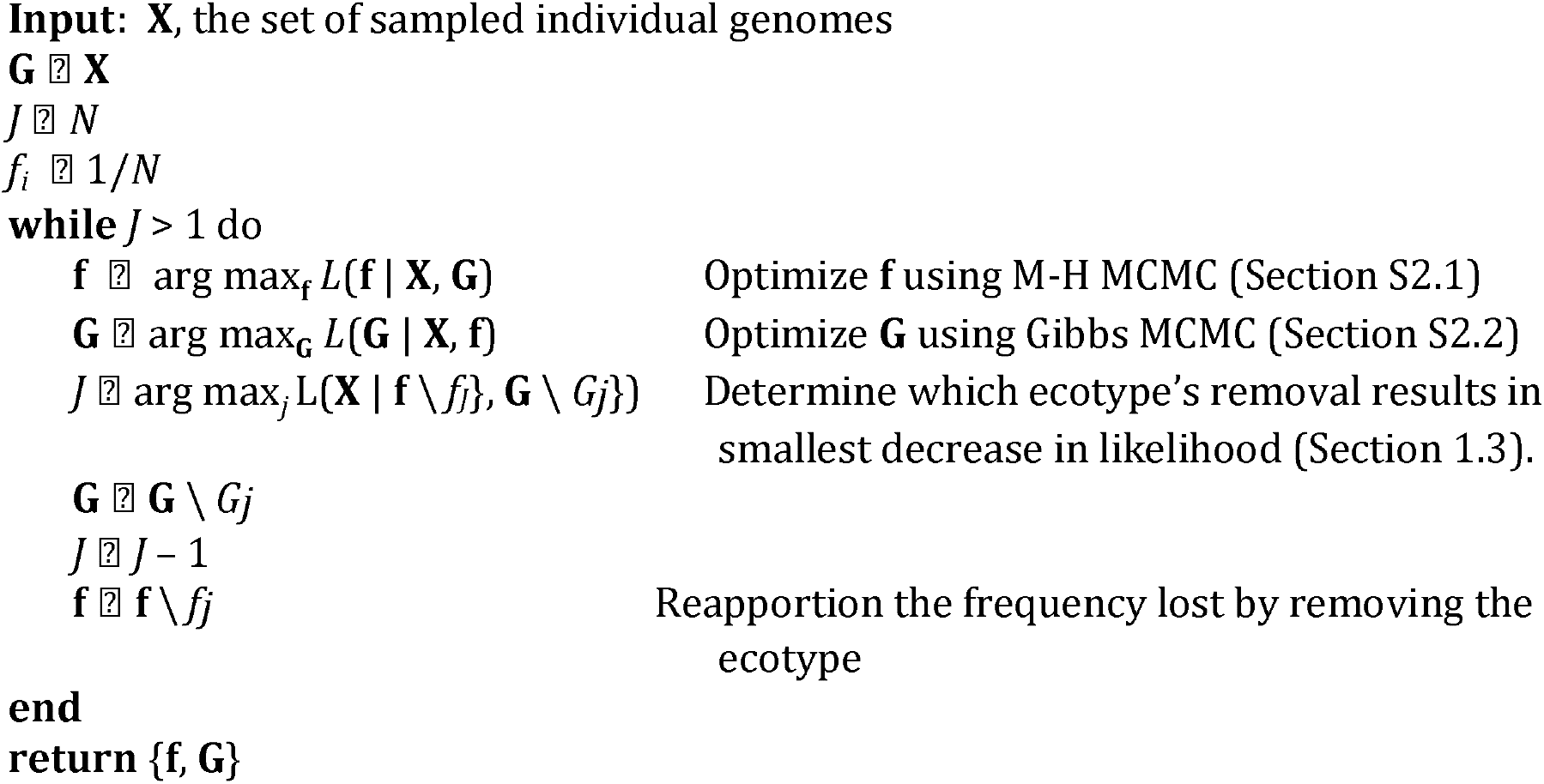

## SI 3: Details of the BPP Methods

Bayesian Phylogenetics and Phylogeography (BPP) is a Bayesian Markov Chain Monte Carlo (MCMC) method for sequence-based species (here, ecotype) delimitation under the multispecies coalescent model (7, 8).

In BPP, the user assigns individuals to populations, the finest feasible division of individuals into ecotypes, and provides a guide tree which serves as a preliminary phylogeny for these populations. BPP may join multiple populations into a single ecotype or call a single population an ecotype, but it will never split a population into multiple ecotypes. BPP has four major categories of analysis, each defined by whether the guide tree and ecotype delimitation are fixed. The analysis we conducted (called A11, or unguided species delimitation) conducts joint ecotype delimitation and ecotype tree inference, meaning that neither is fixed (8). It does this inference through a two-step MCMC algorithm. One step, nearest-neighbor interchange, is used to move between ecotype phylogenies while holding the delimitation constant. The second, reversible-jump MCMC, is used to consider changes to the ecotype delimitations by joining and splitting nodes in the population phylogeny. This process may join two sister populations into a single ecotype, but it will never split a single population into multiple ecotypes. See Yang and Rannala (8) for more details on this method.

We used subsets of the whole genome sequences because use of the entire dataset was computationally prohibitive. We subsetted the data in twelve different ways. Nine of these used the intergenic transcribed spacer sequences (549 bp), one used 10% of all the whole genome sequences (162 kbp), one used 0.1% of all the whole genome sequences (1.6 kbp), and one used the whole genome sequences from only 9 individuals (1.6 Mbp). Details and results of these analyses are in Table S3.1. For all but the analyses of the ITS data, we used the same set of fundamental BPP parameters (Table S3.2). For the analyses of the ITS data, we sought to test the limits of BPP on prokaryotic data. We did this by changing the following: which individuals we included in our analysis (the entire population, only individuals from a single clade, individuals from a few disparate clades, etc.), how we assigned individuals to populations (large populations, every individual in its own population, etc.), and the guide tree (realistic or very scrambled).

Two parameters that we never altered are the priors for {*τ*_*S*_} (the species divergence times expressed in mutations per base) and {*θ*_*S*_}(the average proportion of sites that are different between two randomly selected individuals in a population expressed in substitutions per site). We estimated the expected values of these distributions based on information from Kashtan *et al*. (p. 16 of their Supplemental Materials of ref. (9)). They estimated *θ* = 0.05 from a coalescent simulation of neutral evolution of the largest *Prochlorococcus* ecotype. This *θ* value corresponded to a time to most recent common ancestor of 2.5 × 10^8^ generations, given *μ* = 10 ^−10^ mutations per base per generation. We used this information to estimate the age of the root: *τ* ≈ (2.5 *x* 10^8^ generations) × (10^−10^ mutations / base / generation) = 0.025 mutations / base (9). Consequently, we selected parameters for the inverse-gamma prior for *τ* _*S*_ and *θ* _*S*_ such that E [*τ*_*S*_] = 0.025 and E [*θ*_*S*_] = 0.05. The important BPP parameters are described in Table S3.2.

**Table SI 3.1.**
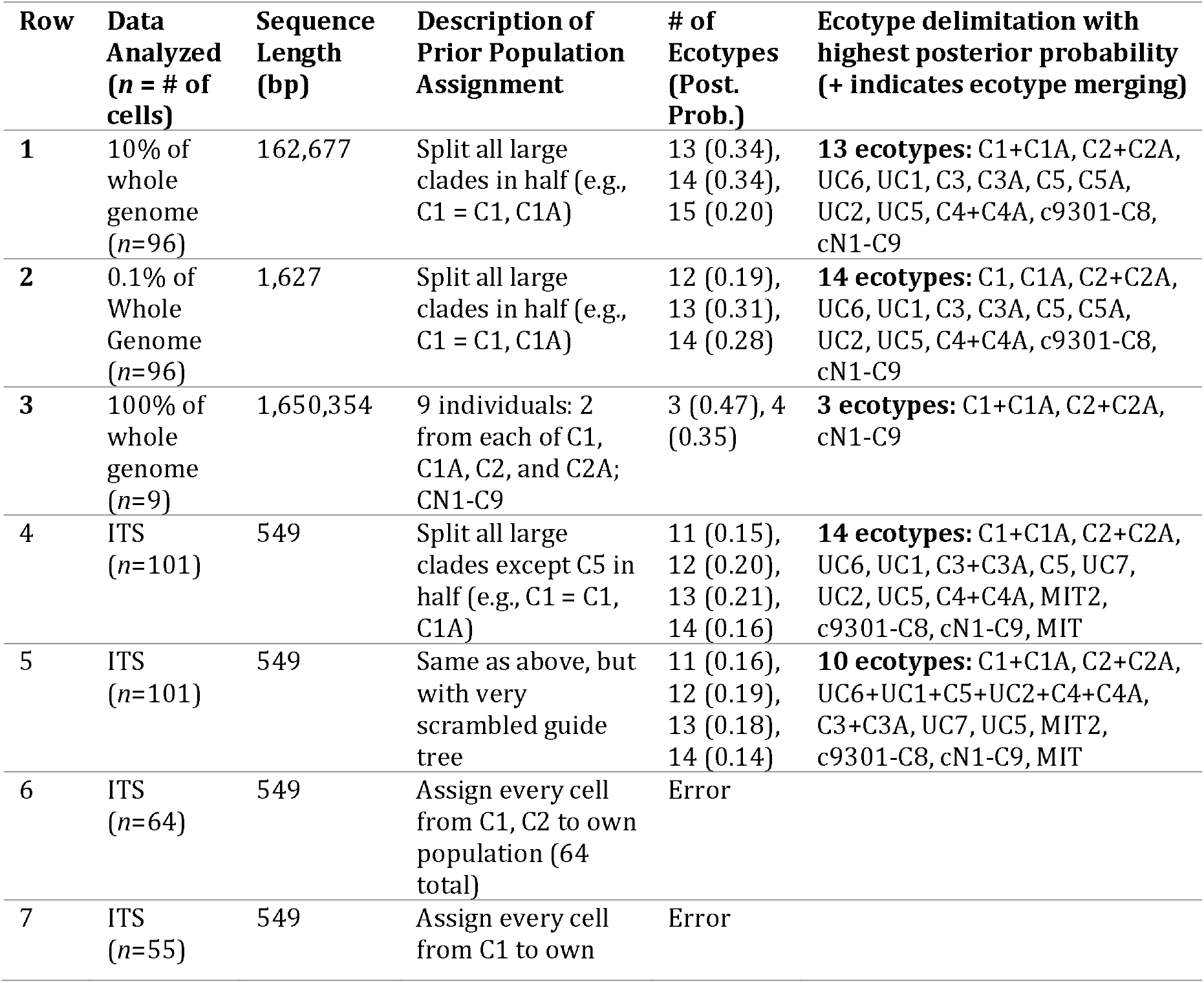
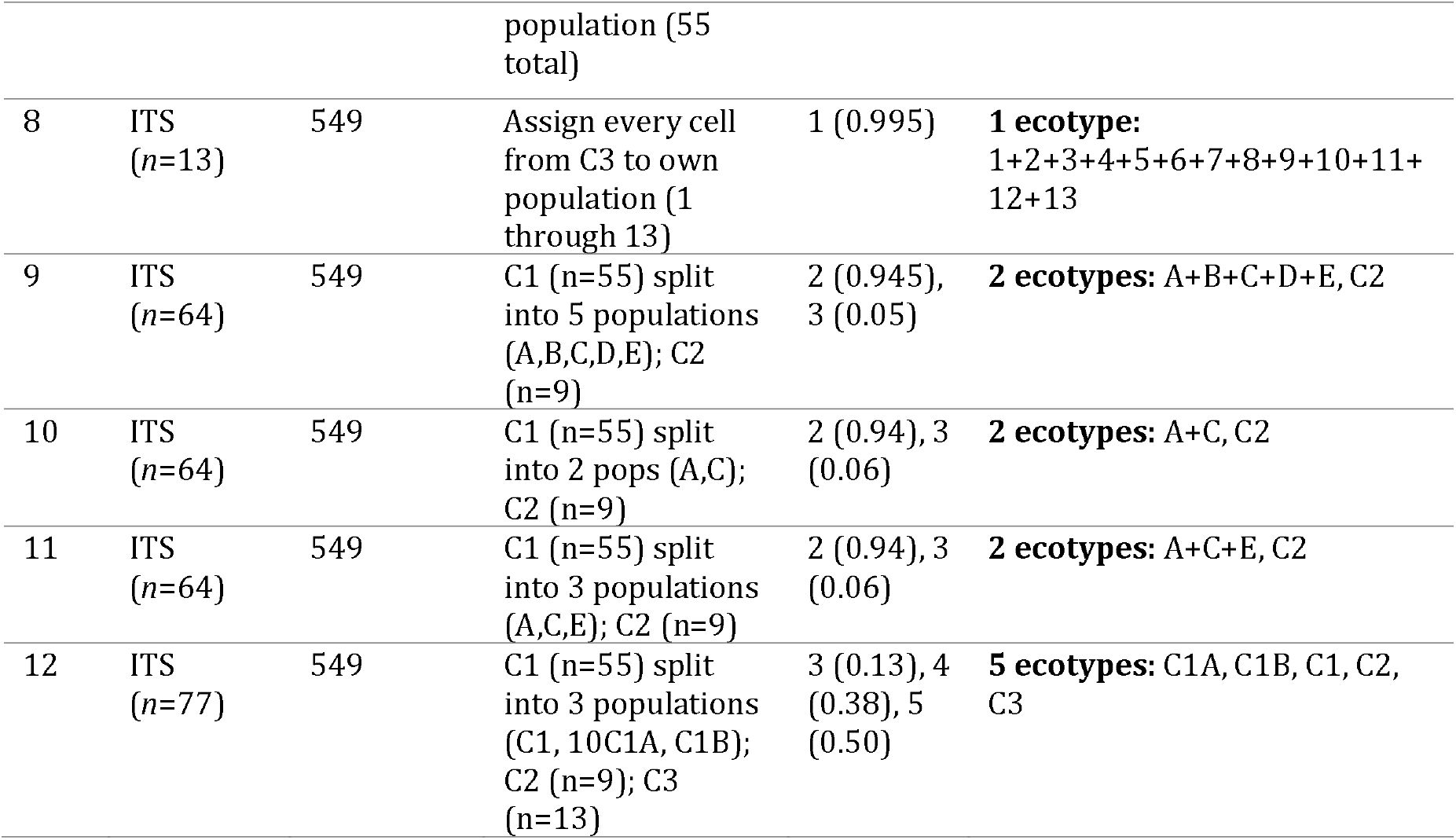
Results of Twelve BPP Analyses. Each row describes a different BPP analysis. The first column shows the data used (ITS data or a subset of the whole genome sequence data) and the number of individual cells included. The second column gives the sequence length for the data analyzed. The third column describes how the cells were allocated into prior populations for the BPP analysis. (Recall that these prior populations can be merged but not subdivided by BPP; see Appendix 3 for more details.) See Figure 2 for a depiction of the guide tree used for all analyses unless otherwise noted and the definitions of the original clades (e.g., C1, c9301-C8, etc.). The final two columns give the results of each analysis: the posterior probability assigned to each number of possible ecotypes, and the ecotype delimitation with the highest posterior probability. The ecotype with the highest posterior probability is indicated as a list of ecotypes, with the plus sign indicating that two prior populations have been merged into one ecotype in the final delimitation. Note that the single ecotype delimitation with the highest posterior probability does not necessarily align with the highest posterior probability number of ecotypes, which accounts for all possible delimitations with a given number of ecotypes. See Figure 3 for the full posterior probability distributions for the analyses in rows 1, 2, and 4.mess

| Row | Data Analyzed (n = # of cells) | Sequence Length (bp) | Description of Prior Population Assignment | # of Ecotypes (Post. Prob.) | Ecotype delimitation with highest posterior probability (+ indicates ecotype merging) |
| --- | --- | --- | --- | --- | --- |
| 1 | 10% of whole genome (n=96) | 162,677 | Split all large clades in half (e.g., C1 = C1, C1A) | 13 (0.34),<br>14 (0.34),<br>15 (0.20) | **13 ecotypes:** C1+C1A, C2+C2A, UC6, UC1, C3, C3A, C5, C5A, UC2, UC5, C4+C4A, c9301-C8, cN1-C9 |
| 2 | 0.1% of Whole Genome (n=96) | 1,627 | Split all large clades in half (e.g., C1 = C1, C1A) | 12 (0.19),<br>13 (0.31),<br>14 (0.28) | **14 ecotypes:** C1, C1A, C2+C2A, UC6, UC1, C3, C3A, C5, C5A, UC2, UC5, C4+C4A, c9301-C8, cN1-C9 |
| 3 | 100% of whole genome (n=9) | 1,650,354 | 9 individuals: 2 from each of C1, C1A, C2, and C2A; CN1-C9 | 3 (0.47), 4 (0.35) | **3 ecotypes:** C1+C1A, C2+C2A, cN1-C9 |
| 4 | ITS (n=101) | 549 | Split all large clades except C5 in half (e.g., C1 = C1, C1A) | 11 (0.15),<br>12 (0.20),<br>13 (0.21),<br>14 (0.16) | **14 ecotypes:** C1+C1A, C2+C2A, UC6, UC1, C3+C3A, C5, UC7, UC2, UC5, C4+C4A, MIT2, c9301-C8, cN1-C9, MIT |
| 5 | ITS (n=101) | 549 | Same as above, but with very scrambled guide tree | 11 (0.16),<br>12 (0.19),<br>13 (0.18),<br>14 (0.14) | **10 ecotypes:** C1+C1A, C2+C2A, UC6+UC1+C5+UC2+C4+C4A, C3+C3A, UC7, UC5, MIT2, c9301-C8, cN1-C9, MIT |
| 6 | ITS (n=64) | 549 | Assign every cell from C1, C2 to own population (64 total) | Error |  |
| 7 | ITS (n=55) | 549 | Assign every cell from C1 to own population (55 total) | Error |  |
| 8 | ITS<br>(n=13) | 549 | Assign every cell from C3 to own population (1 through 13) | 1 (0.995) | **1 ecotype:**<br>1+2+3+4+5+6+7+8+9+10+11+12+13 |
| 9 | ITS<br>(n=64) | 549 | C1 (n=55) split into 5 populations (A,B,C,D,E); C2 (n=9) | 2 (0.945),<br>3 (0.05) | **2 ecotypes:** A+B+C+D+E, C2 |
| 10 | ITS<br>(n=64) | 549 | C1 (n=55) split into 2 pops (A,C); C2 (n=9) | 2 (0.94), 3<br>(0.06) | **2 ecotypes:** A+C, C2 |
| 11 | ITS<br>(n=64) | 549 | C1 (n=55) split into 3 populations (A,C,E); C2 (n=9) | 2 (0.94), 3<br>(0.06) | **2 ecotypes:** A+C+E, C2 |
| 12 | ITS<br>(n=77) | 549 | C1 (n=55) split into 3 populations (C1, 10C1A, C1B); C2 (n=9); C3 (n=13) | 3 (0.13), 4<br>(0.38), 5<br>(0.50) | **5 ecotypes:** C1A, C1B, C1, C2, C3 |

**Table SI 3.2.** BPP Parameters. Typical parameter values used to run BPP analyses. Note that the mean of θ and τ_s (root age) prior distributions were informed by the neutral coalescent simulation run by Kashtan et al. (9) which sought to mimic characteristics of *Prochlorococcus* (section 6.2 of the Supplemental Information of ref. (9)). E[θ] = 0.05, E[τ_s] = 0.025 mutations/base.

| Code in BPP Ctl File | Meaning |
| --- | --- |
| Speciesdelimitation = 1 1 2 1 | First “1” means species assignments are not given by the user. Subsequent values specify rjMCMC algorithm and parameters. |
| Speciestree = 1 0.4 0.2 0.1 | First “1” means the given species tree is used as the guide tree in the rjMCMC run for species delimitation. Subsequent values are parameters. |
| Speciesmodelprior = 3 | Each number of species is assigned an equal prior probability; probability divided uniformly among compatible models of species delimitation. (Best choice when many populations; avoids biasing towards many species) |
| Cleandata = 0 | Includes columns with ambiguity data in the likelihood calculation |
| Thetaprior = 3 0.1<br>($\theta \sim \text{IG}(3, 0.1)$, $E[\theta]=0.05$) | Theta parameters estimated (rather than being integrated out using conjugate prior) according to an inverse gamma prior |
| Tauprior = 3 0.05<br>($\tau_s \sim \text{IG}(3, 0.05)$, $E[\tau_s]=0.025$) | Specifies inverse gamma prior for $\tau_s$, the divergence time parameter for the root in the species tree. |
| finetune = 1: .01 .02 .03 .04 .05 .01 .01 | First “1” specifies to automatically adjust MCMC step lengths. Subsequent values are initial step lengths for various parameters. |

## SI 4: Results of *TreeFree*

The sample IDs and clade assignment of *Prochlorococcus* genomes from Kashtan *et al*. (9). The last two columns show the ecotype to which *TreeFree* assigned each genome with highest posterior probability based on either 10% or 0.1% of the sequences.

| Sample ID | Clade | 10% | 0.1% |
| --- | --- | --- | --- |
| 526B17 | C1 | 4 | 0 |
| 526B19 | UC | 4 | 0 |
| 526B22 | C1 | 4 | 0 |
| 526D20 | C1 | 4 | 0 |
| 526K3 | C1 | 4 | 0 |
| 526N5 | C5 | 4 | 0 |
| 526N9 | C1 | 4 | 0 |
| 527E14 | C2 | 22 | 0 |
| 527E15 | C3 | 9 | 0 |
| 527G5 | C1 | 4 | 0 |
| 527I9 | C1 | 0 | 0 |
| 527L15 | C1 | 4 | 0 |
| 527L16 | C1 | 20 | 0 |
| 527L22 | c9301 | 10 | 2 |
| 527N11 | C1 | 1 | 0 |
| 527P5 | C1 | 23 | 0 |
| 528J14 | C2 | 22 | 0 |
| 528J8 | cN1 | 8 | 0 |
| 528K19 | C1 | 1 | 0 |
| 528N17 | C4 | 11 | 0 |
| 528N20 | C1 | 13 | 0 |
| 528N8 | C1 | 18 | 0 |
| 528O2 | UC | 4 | 6 |
| 528P14 | c9301 | 10 | 2 |
| 528P18 | C3 | 2 | 5 |
| 529B19 | C1 | 16 | 0 |
| 529C4 | C1 | 3 | 0 |
| 529D18 | C3 | 2 | 5 |
| 529J11 | C1 | 4 | 0 |
| 529J15 | C3 | 9 | 5 |
| 529J16 | C1 | 4 | 0 |
| 529O19 | C1 | 0 | 0 |
| 495D8 | UC | 4 | 0 |
| 495G23 | UC | 4 | 0 |
| 495I8 | C1 | 4 | 0 |
| 495K23 | C1 | 7 | 0 |
| 495L20 | C3 | 2 | 5 |
| 495N16 | C1 | 20 | 0 |
| 495N3 | C4 | 5 | 0 |
| 495N4 | C1 | 18 | 0 |
| 495P20 | UC | 6 | 0 |
| 496A2 | C3 | 2 | 5 |
| 496E10 | C1 | 7 | 0 |
| 496G15 | C2 | 4 | 0 |
| 496M6 | UC | 4 | 0 |
| 496N4 | C1 | 13 | 0 |
| 497E17 | cN1 | 8 | 0 |
| 497I20 | C1 | 4 | 0 |
| 497J18 | C3 | 9 | 5 |
| 497N18 | UC | 22 | 0 |
| 498A3 | c9301 | 10 | 3 |
| 498B22 | C2 | 22 | 0 |
| 498B23 | C4 | 11 | 0 |
| 498C16 | C2 | 22 | 0 |
| 498F21 | C1 | 12 | 0 |
| 498G3 | C1 | 4 | 0 |
| 498I20 | C5 | 4 | 1 |
| 498J20 | C1 | 4 | 0 |
| 498L10 | C1 | 13 | 0 |
| 498M14 | C1 | 4 | 0 |
| 498N4 | C3 | 2 | 5 |
| 498N8 | C2 | 22 | 0 |
| 498P15 | C1 | 14 | 0 |
| 498P3 | C1 | 16 | 0 |
| 518A17 | C3 | 2 | 5 |
| 518A6 | C3 | 2 | 5 |
| 518D8 | C1 | 15 | 0 |
| 518E10 | C1 | 4 | 0 |
| 518I6 | C5 | 4 | 4 |
| 518J7 | C3 | 2 | 5 |
| 518K17 | C1 | 16 | 0 |
| 518O7 | C1 | 4 | 0 |
| 519A13 | UC | 4 | 0 |
| 519B7 | C4 | 5 | 0 |
| 519C7 | C1 | 17 | 0 |
| 519D13 | C1 | 1 | 0 |
| 519E23 | C1 | 21 | 0 |
| 519G16 | C3 | 2 | 5 |
| 519L21 | C1 | 4 | 0 |
| 519O11 | C1 | 20 | 0 |
| 519O21 | c9301 | 10 | 2 |
| 520B18 | C1 | 0 | 0 |
| 520D2 | C1 | 19 | 0 |
| 520E22 | C1 | 7 | 0 |
| 520F22 | C2 | 22 | 0 |
| 520K10 | cN1 | 8 | 0 |
| 520M11 | C2 | 22 | 0 |
| 521A19 | C1 | 1 | 0 |
| 521B10 | C1 | 23 | 0 |
| 521C8 | C3 | 2 | 0 |
| 521K15 | C1 | 21 | 0 |
| 521M10 | C1 | 16 | 0 |
| 521N3 | C1 | 18 | 0 |
| 521N5 | C1 | 4 | 0 |
| 521O20 | C1 | 20 | 0 |
| 521O23 | C1 | 0 | 0 |

## Notes

### Competing Interest Statement

The authors have declared no competing interest.

